# Defining tertiary sulci in lateral prefrontal cortex in chimpanzees using human predictions

**DOI:** 10.1101/2022.04.12.488091

**Authors:** Catherine B. Hathaway, Willa I. Voorhies, Neha Sathishkumar, Chahat Mittal, Jewelia K. Yao, Jacob A. Miller, Benjamin J. Parker, Kevin S. Weiner

**Author notes:** Co-first author (Catherine B. Hathaway, Willa I. Voorhies). Corresponding author: Kevin S. Weiner.

## Abstract

Similarities and differences in brain structure and function across species is of major interest in systems neuroscience, comparative biology, and brain mapping. Recently, increased emphasis has been placed on tertiary sulci, which are shallow indentations of the cerebral cortex that appear last in gestation, continue to develop after birth, and are largely either human- or hominoid-specific. While tertiary sulcal morphology in lateral prefrontal cortex (LPFC) has been linked to functional representations and cognition in humans, it is presently unknown if LPFC tertiary sulci also exist in non-human hominoids. To fill this gap in knowledge, we leveraged two freely available multimodal datasets to address the following main question: Can LPFC tertiary sulci be defined in chimpanzee cortical surfaces from human predictions? We found that 1-3 components of the posterior middle frontal sulcus (pmfs) in the posterior middle frontal gyrus are identifiable in nearly all chimpanzee hemispheres. In stark contrast to the consistency of the pmfs components, we could only identify components of the paraintermediate frontal sulcus (pimfs) in two chimpanzee hemispheres. LPFC tertiary sulci were relatively smaller and shallower in chimpanzees compared to humans. In both species, two of the pmfs components were deeper in the right compared to the left hemisphere. As these results have direct implications for future studies interested in the functional and cognitive role of LPFC tertiary sulci across species, we share probabilistic predictions of the three pmfs components to guide the definitions of these sulci in future studies.

## INTRODUCTION

Similarities and differences in brain structure and function across species is of major interest in systems neuroscience and comparative biology. Recently, increased emphasis has been placed on tertiary sulci, which are shallow indentations of the cerebral cortex that appear late in gestation, continue to develop after birth, and are related to the organization of cortical networks (Connolly 1950; Welker 1990; Armstrong et al. 1995; Weiner 2019; Lopez-Persem et al. 2019; Miller et al. 2020; Miller et al. 2021a, 2021b). Additionally, the morphology of some tertiary sulci is related to cognition and behavior (Amiez et al. 2018; Voorhies et al. 2021) with translational and clinical applications (Garrison et al. 2015; Ammons et al. 2021). Tertiary sulci are present in hominoid brains, but not other widely studied animals in neuroscience research such as mice, marmosets, and macaques (Amiez et al.; Lopez-Persem et al. 2019; Miller et al. 2020; Voorhies et al. 2021; Miller et al. 2020; Miller et al. 2021a, 2021b).

Intriguingly, some tertiary sulci are human-specific, while other tertiary sulci are only present in some, but not all human brains. For example, the mid-fusiform sulcus in ventral temporal cortex and the inframarginal sulcus in posterior cingulate cortex are present in every human brain (Miller et al. 2020; Willbrand et al. 2021), while the paracingulate sulcus in medial frontal cortex (Paus et al. 1996; Fornito et al. 2004, 2006, 2008; Amiez et al. 2018) and the paraintermediate frontal sulcus in lateral prefrontal cortex (Amiez and Petrides 2007; Voorhies et al. 2021) are not. The former three sulci also have been studied in non-human hominoid brains such as chimpanzees (Amiez et al., 2019; Amiez et al. 2021; Miller et al. 2020; Willbrand et al. 2021), while the latter has not. To fill this gap in knowledge, we focus on tertiary sulci in lateral prefrontal cortex (LPFC) in the present study and ask two main questions: 1) Can LPFC tertiary sulci be defined in chimpanzee cortical surfaces from human predictions? and 2) As surface area and depth are defining features differentiating tertiary sulci from primary and secondary sulci (Connolly 1950; Welker 1990; Armstrong et al. 1995; Weiner 2019; Lopez-Persem et al. 2019; Miller et al. 2020; Miller et al. 2021a, 2021b), do LPFC tertiary sulci in chimpanzees differ in the relative surface area and relative depth compared to LPFC tertiary sulci identified in humans?

## RESULTS

To answer these questions, we leveraged two freely available multimodal datasets: The National Chimpanzee Brain Resource (https://www.chimpanzeebrain.org/) and The Human Connectome Project (http://www.humanconnectomeproject.org/). Briefly, cortical surface reconstructions were generated for both species from T1 images using FreeSurfer (https://www.freesurfer.net). Leveraging our previously published pipeline that accurately projects probabilistic definitions of sulci defined in the human cerebral cortex to individual chimpanzee hemispheres (Miller et al., 2020), we tested if LPFC tertiary sulci could be defined in chimpanzee cortical surfaces from human predictions (Figure 1). If possible, we then used our previously published morphological pipeline to statistically test if relative surface area and relative depth of LPFC tertiary sulci differed between humans and chimpanzees. For comparison, we also included surrounding sulci, though the main focus was on tertiary sulci. We report five main findings.

**Figure 1.**
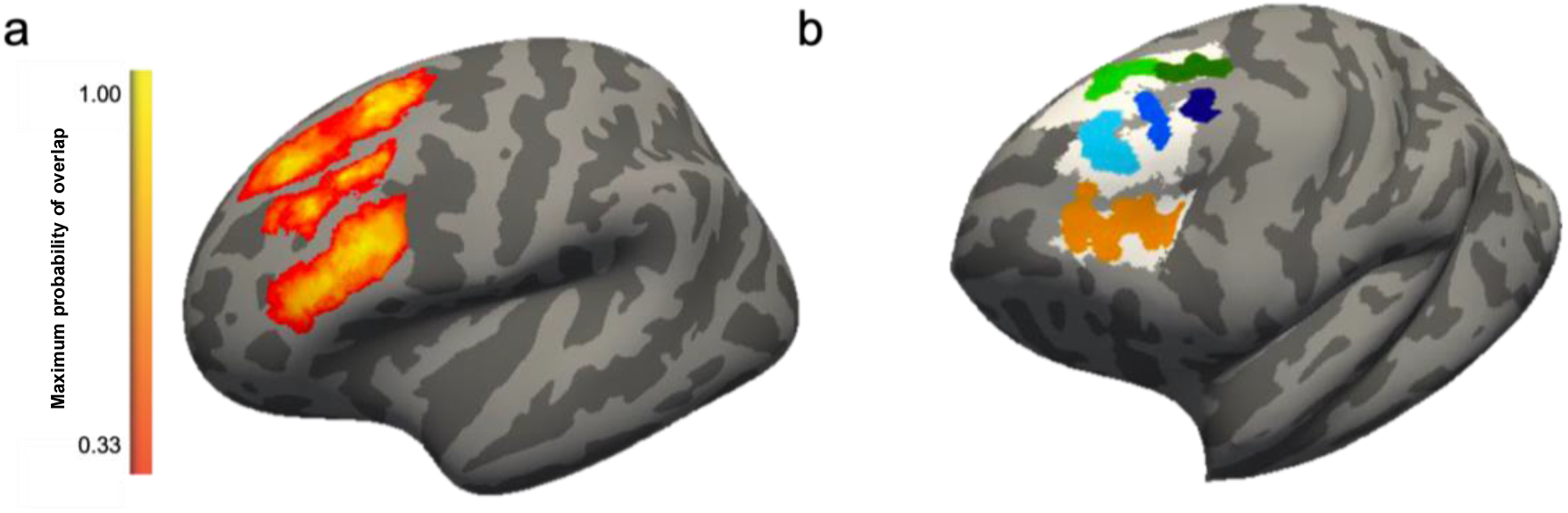
Manual labeling protocol of LPFC tertiary sulci in chimpanzees guided by human predictions. a) *fsaverage* surface showing maximum sulcal probability maps of LPFC sulci from previous work (Miller et al., 2021a). Binarized maps were used to guide labeling on chimpanzee cortical surfaces. Sulcal maps were thresholded at 33% to minimize overlap for visualization purposes. b) Example chimpanzee inflated cortical surface illustrating manual labeling procedure. Binarized maximum probability maps in humans were projected to individual chimpanzee cortical surfaces (*white*). These projections were used to guide manual sulcal labeling (colors) for the ifs (orange), sfs-p (dark green), sfs-a (light green), pmfs-p (dark blue), pmfs-i (blue), and pmfs-a (light blue) in individual chimpanzee hemispheres.

First, some, but not all, LPFC tertiary sulci are consistently identifiable in chimpanzees. Specifically, tertiary pmfs components were identifiable in a majority, but not all, chimpanzee hemispheres (lh: 83%; rh: 79%; Figure 2b; Supplementary Figure 2a). Interestingly, the incidence rates of pmfs components in chimpanzees were significantly less than in humans (lh: *χ**^2^*** = 5.54, *p* = 0.01; rh: *χ**^2^*** = 11.7, *p* < 0.001), as humans consistently had three identifiable pmfs components in every hemisphere, but chimpanzees did not. Additionally, while at least one pimfs component was identifiable nearly 100% of the time in the human brain, a pimfs component was only identifiable in 2 of the 60 chimpanzee hemispheres measured (Figure 2a, Supplementary Figure 1). Due to the consistent identification of the pmfs components in chimpanzees, and the consistent absence of the pimfs components in chimpanzees, we focus our cross-species morphological analyses on the pmfs components. We emphasize for the reader that though previous studies have identified a middle frontal sulcus in the brains and endocasts of chimpanzees (Bailey et al. 1950; Connolly 1950; Sherwood et al. 2003; Schenker et al. 2010; Falk et al. 2018), these sulcal definitions are often distinct from tertiary pmfs definitions as discussed previously (Miller et al., 2021a, 2021b).

**Figure 2.**
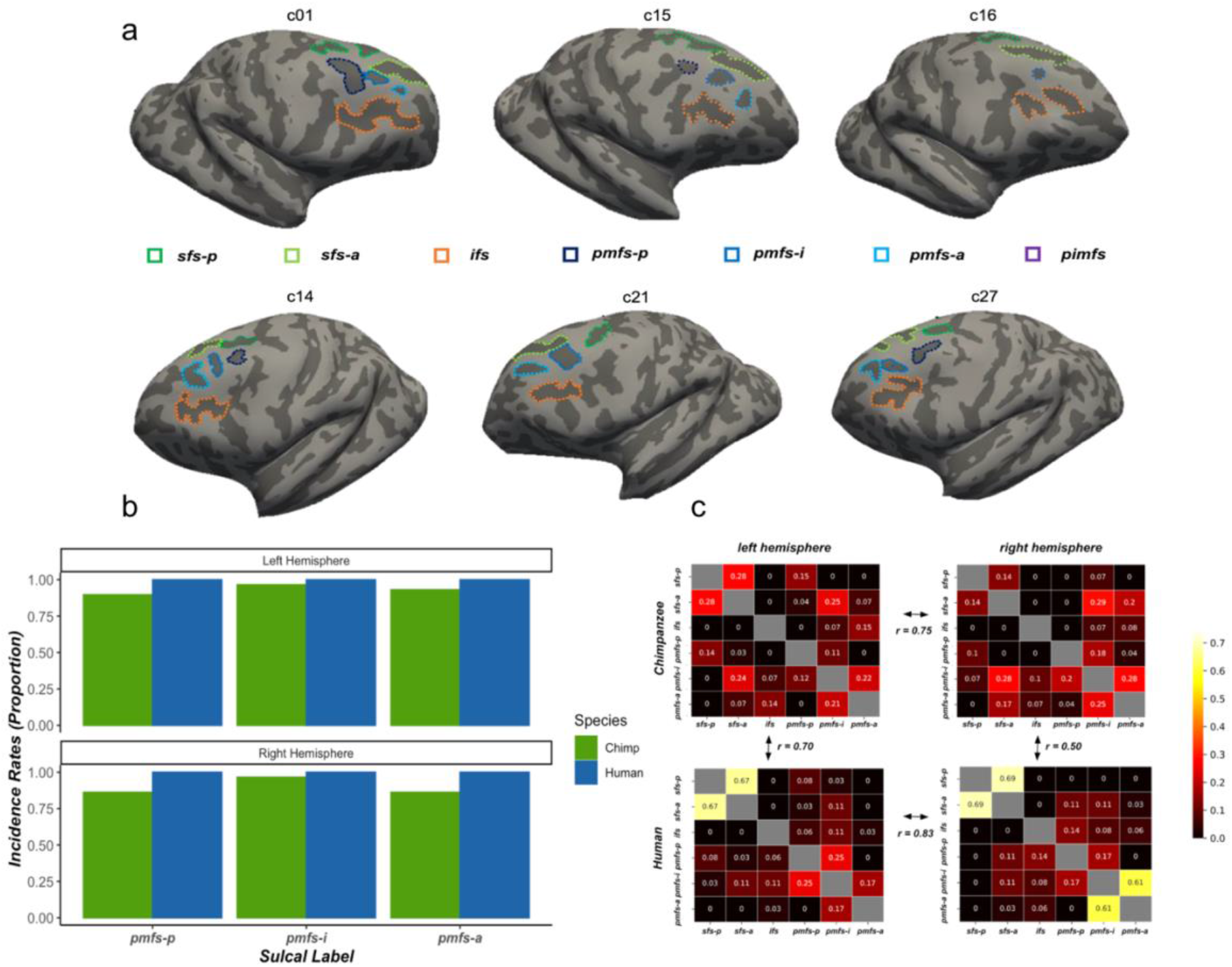
Tertiary sulci in lateral prefrontal cortex are identifiable in chimpanzees using human sulcal predictions. a) Example right (*top*) and left (*bottom*) sulcal labels for 6 chimpanzees. Chimpanzees had between 1 and 4 identifiable tertiary sulci in a given hemisphere. b) Comparison of tertiary sulcal incidence rates (pmfs components) for chimpanzees and humans. Incidence rates of pmfs components in chimpanzees were significantly less than in humans (lh: *χ**^2^*** = 5.54, *p* = 0.01; rh: *χ**^2^*** = 11.7, *p* < 0.001). c) For each sulcus, we report the proportion of intersection (frequency of occurrence/total number of observations in the hemisphere) with every other LPFC sulcus included in the present study (see colorbar for reference; empty white cells in the matrix reflect the fact that a sulcus cannot intersect with itself). ifs: inferior frontal sulcus; pimfs: paraintermediate frontal sulcus; pmfs-a, pmfs-i, pmfs-p: anterior, intermediate, and posterior components of the posterior middle frontal sulcus; sfs-a, sfs-p: anterior and posterior components of the superior frontal sulcus. Similarity between hemispheres and species are reported as Pearson’s r correlation coefficients.

Second, quantifying tertiary sulcal variability by examining the prevalence of sulcal types (Figure 2c) based on their rate of intersection with neighboring sulci (Materials and Methods) reveals similar rates of intersections between left and right hemispheres in chimpanzees (Pearson’s r = 0.75) and humans (Pearson’s r = 0.83). In general, humans and chimpanzees showed similar patterns of sulcal intersections (Figure 2c) with higher similarity in the left compared to the right hemisphere (lh: Pearson’s r = 0.70; rh: Pearson’s r = 0.50). In the left hemisphere, the posterior and anterior portions of the sfs showed higher rates of intersection in humans (Figure 2c), while in the right hemisphere, humans showed higher rates of intersection between intermediate and anterior pmfs components (Figure 2c).

Third, for sulcal depth, a mixed effects linear model with sulcus, hemisphere, and species as factors showed that (i) sulci were shallower in chimpanzees relative to humans (F(1,63) = 199.32, p < 0.0001, Figure 3a), (ii) depth varied by sulcus (F(5,615) = 86.50, p < 0.0001), and (iii) in both species, the pmfs components were shallower (*Mean*(*sd*): pmfs-p = 10.4(4.73), pmfs-i = 10.8(4.39), pmfs-a = 11.2(4.29)) than the surrounding ifs and sfs (*Mean*(*sd*): ifs = 17.0(2.39), sfs-p = 14.4(3.28), sfs-a = 12.8(3.44)) components (Figure 3a; Supplementary Figure 2b for raw depth values). There was also a main effect of hemisphere in which sulci in the left hemisphere were shallower than the right hemisphere in both species (F(1,63) = 11.82, p = 0.001; *left*: Chimpanzee mean(sd) = 10.5(4.46), Human mean(sd) = 13.9(4.07); *right*: Chimpanzee mean(sd) =10.9(4.11), Human mean(sd) = 15.0(3.63). Post-hoc analyses revealed that the differences between species were most pronounced for the tertiary sulci and the sfs components (Figure 3a; all *p*s < 0.0001).

**Figure 3.**
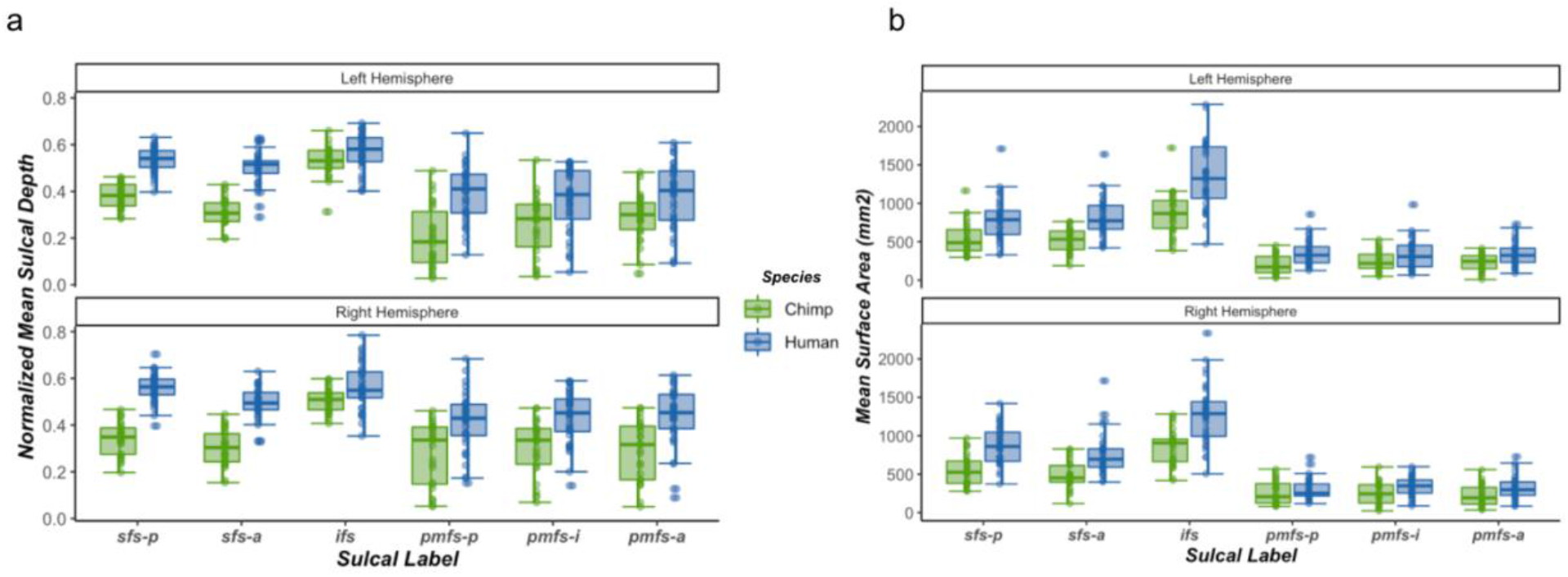
LPFC sulci are relatively smaller and shallower in chimpanzees compared to humans. a) Normalized mean sulcal depth (mm) for each sulcus in chimpanzees (green) and humans (blue) for the left and right hemispheres. Sulci were consistently shallower in chimpanzees. Normed sulcal depth is calculated as a proportion relative to the deepest point in the hemisphere. b) The same as a) but for relative mean surface area (mm^2^). Sulci were relatively larger in humans than chimpanzees. Horizontal lines represent median values, boxes represent interquartile range, and whisker lines represent the 1st and 3rd quartiles.

Fourth, for surface area, a mixed effects linear model with the same three factors showed an expected species effect in which all sulci were less prominent in chimpanzees than in humans (F(1,63) = 122.47, p < 0.0001; *left*: Chimpanzee mean(sd) = 438.05(296.81), Human mean(sd) = 665.25(450.51); *right*: Chimpanzee mean(sd) = 448.68(282.61), Human mean(sd) = 643.85(430.22). Post-hoc analyses further showed that differences were most pronounced for non-tertiary sulci (*p*s< 0.001; Figure 3b). There was also an effect of sulcal type (F(5,615) = 307.09, p < 0.0001) in which the pmfs components (*Mean*(*sd*): pmfs-p = 286(155), pmfs-i = 300(156), pmfs-a = 285(150)) were smaller than the ifs and sfs *Mean*(*sd*): (ifs = 1109(411), sfs-p = 705(275), sfs-a = 658(264)) components across species and hemispheres (Figure 3b).

Fifth, we replicated and extend previous findings showing that the ifs was comparably deep between the two hemispheres in chimpanzees with little asymmetry (Bogart et al. 2012). We also find that the ifs does not show hemispheric asymmetry in either species and report a comparable asymmetry value in chimpanzees (*chimpanzee*: mean(sd) = −0.02 (0.07); *human*: mean(sd) = −0.0002(0.07)) as previously reported. We also extend these previous results by considering tertiary sulci and show that, in both chimpanzees and humans, the pmfs-i and pmfs-p showed significant rightward asymmetry (*pmfs-p*: mean(sd) = 0.18(0.54); *pmfs-i*: mean(sd) = 0.25 (0.58)); Figure 4). Although this effect was present in both species, we did observe a significant species difference in the pmfs-p, in which chimpanzees showed a greater rightward asymmetry than humans (*Mean*(*sd*): chimpanzee = 0.36(0.72) human = 0.07(0.36); *p* = 0.01).

**Figure 4.**
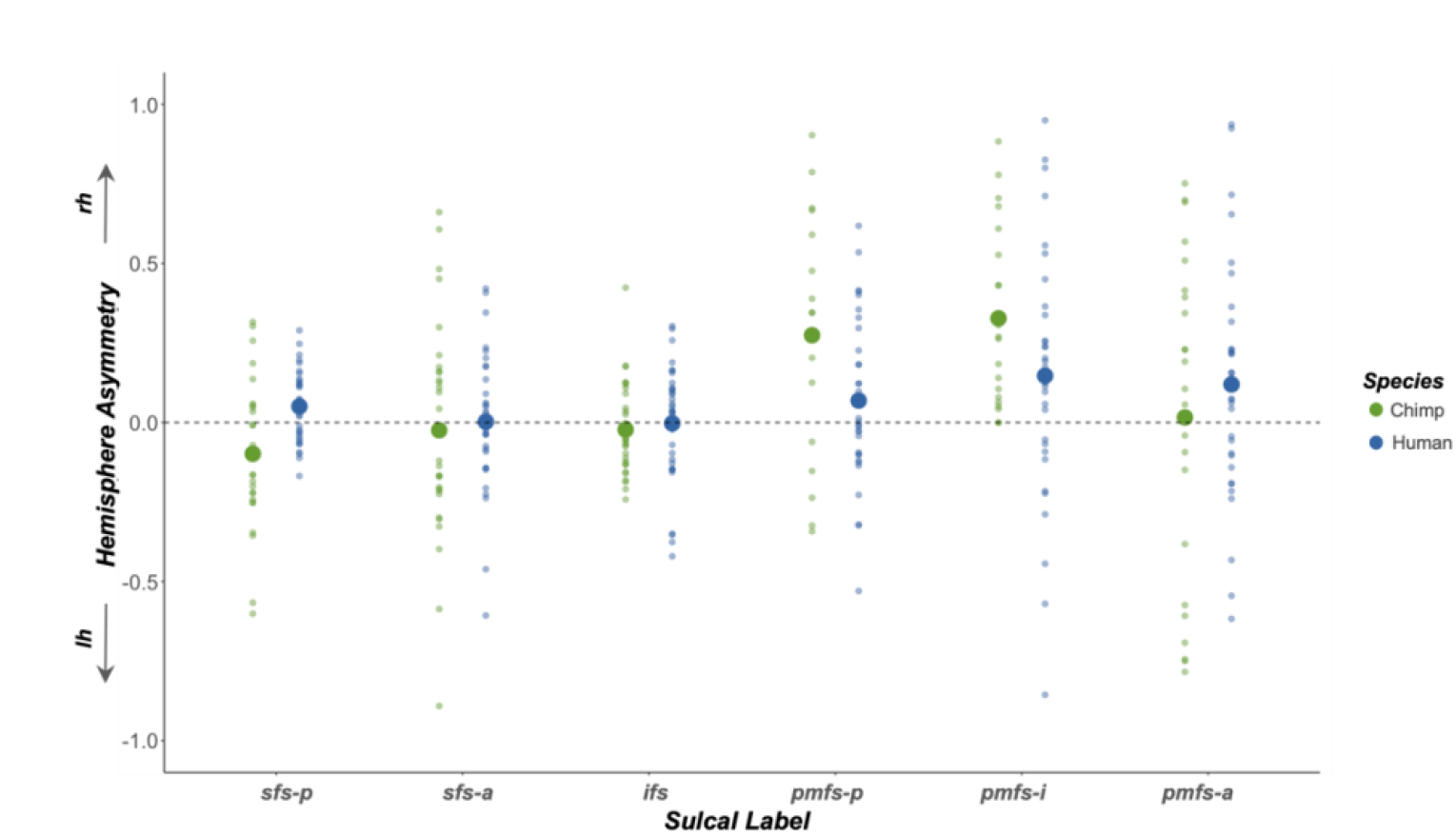
Some LPFC tertiary sulci are deeper in the right compared to the left hemisphere in both chimpanzees and humans. Depth asymmetry for each sulcus in chimpanzees (*green*) and humans (*blue*). Asymmetry was calculated as (RH - LH)/(RH + LH)*2. Large dots represent mean asymmetry values across species. Small dots reflect asymmetry for individual members in each species. Deeper sulci in the right hemisphere are above zero, while deeper sulci in the left hemisphere are below zero.

## DISCUSSION

To our knowledge, the present findings are the first to identify and quantify morphological features of the three shallow components of the posterior middle frontal sulcus (pmfs-a, pmfs-i, and pmfs-p) in LPFC of chimpanzees. In a recent historical analysis and review of the literature (Miller et al., 2021b), the pmfs components were largely overlooked in previous studies due to their variability in both human and non-humanoid primate brains. Nevertheless, previous studies often mentioned the presence and variability of sulcal components within the posterior MFG of chimpanzees (Bailey et al. 1950; Connolly 1950; Sherwood et al. 2003; Schenker et al. 2010; Falk et al. 2018). In direct reference to this variability, Falk and colleagues (2018) write:

“These newly identified configurations for *fm* show that variation in chimpanzee frontal lobes includes more complex midfrontal gyri than previously described [Connolly 1950; Falk 2014].”

Here, we explicitly quantify that these “newly identified configurations” of the pmfs components are more humanlike in their appearance within the chimpanzee LPFC than previously thought, while the pimfs components in the anterior MFG are more rare in chimpanzees than in humans. Interestingly, the presence or absence of the pimfs has been linked to higher-level aspects of cognition in humans (Willbrand et al., 2022), which begs the question: What is the functional and cognitive role of the pimfs in chimpanzees?

Additionally, the present findings provide novel insight into the morphological asymmetry of LPFC tertiary sulci in chimpanzees for the first time. After replicating previous findings of a lack of asymmetry in the depth of the inferior frontal sulcus in chimpanzees (Bogart et al. 2012), we further showed that there is a larger rightward depth asymmetry for an LPFC tertiary sulcus (pms-p) in chimpanzees compared to humans. Thus, future studies can further explore the functional and cognitive meaning of this asymmetry guided by our shared probabilistic predictions of LPFC sulci (Figure 5), as well as further build from the recent foundation showing the similarities and differences in tertiary sulci across the hominoid clade in ventral temporal (Miller et al. 2020), medial prefrontal (Amiez et al., 2019), posterior cingulate (Willbrand et al. 2021), and now lateral prefrontal cortices.

**Figure 5.**
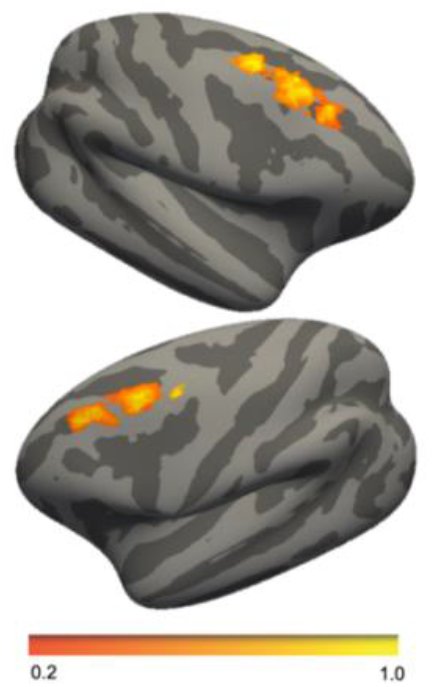
LPFC tertiary sulcal probability maps in chimpanzees. Maximum probability maps for the three consistently identifiable tertiary sulcal labels (*pmfs-p, pmfs-i, pmfs-a*). To generate the maps, each label was transformed from each individual to a custom average template created from 30 additional chimpanzees not included in the original analysis. For each vertex, we calculated the proportion of chimpanzees for whom that vertex is labeled as the given sulcus (the warmer the color, the higher the overlap in each image). In the case of multiple labels for one vertex, the sulcus with the highest overlap across participants was assigned to a given vertex. To reduce spatial overlap for visualization purposes, these maps were thresholded to 20% overlap across chimpanzees.

The present study would not have been possible without using freely available multimodal atlases. The fact that some LPFC tertiary sulci are identifiable in both humans and chimpanzees informs a “horizontal translation” of relating neuroanatomical structures between species. Specifically, our findings show that tertiary sulci within the posterior MFG are consistent between humans and chimpanzees and may serve as a foundation from which to formally compare functional and neuroanatomical features across spatial scales between species. For instance, do certain cytoarchitectonic or functional areas co-localize with these tertiary sulci between species or do they identify transitions in one species, but not another? Additionally, do these sulci have consistent relationships with underlying short or long-range white matter tracts between species? As resting state data are available for both species (Amiez et al. 2021), these sulci can also serve as seeds in functional connectivity analyses in future studies. Interestingly, previous research indicates that the presence or absence of tertiary sulci in medial PFC affects the organization of functional networks – both in the location of the hub of the default mode network as well as the appearance or absence of new clusters elsewhere in the brain, respectively (Lopez-Persem et al. 2019). Thus, the presence or absence of the pimfs components in the human brain may be reflective of individual differences in functional networks within species, while the absence of the pimfs components in chimpanzees may be reflective of differences in functional connectivity across species, which can be tested in future research. Crucially, this prediction in direct relation to the pimfs would not have been generated without these freely available datasets.

In conclusion, our study builds on recent studies showing that tertiary sulci are not just a feature of the human cerebral cortex, but also are commonly identifiable in other hominoid brains. Nevertheless, not all tertiary sulci are frequently identifiable either across human individuals (such as the paracingulate sulcus from previous studies) or across species (such as the paraintermediate frontal sulcus from the present study). Thus, an important goal for future studies is to pinpoint which tertiary sulci are – and which tertiary sulci are not – identifiable in the cerebral cortices within and across species. Here, we show that a useful approach to achieve this goal is to use the definitions of tertiary sulci from human brains to guide the definitions of tertiary sulci in the brains of non-human hominoids. Future research will show the generalizability or specificity of this approach in other cortical expanses and species, as well as functional and cognitive insights that it may provide for understanding the evolution of association cortices, functional representations, and cognition.

## Acknowledgments

This research was supported by an NSF-GRFP fellowship (WIV), a T32 HWNI training grant (BJP), a NICHD R21HD100858 (KSW, SAB), and an NSF CAREER Award 392 2042251 (KSW). Data were provided by the Human Connectome Project, WU-Minn Consortium (Principal Investigators: David Van Essen and Kamil Ugurbil; 1U54MH091657) funded by the 16 NIH Institutes and Centers that support the NIH Blueprint for Neuroscience Research; and by the McDonnell Center for Systems Neuroscience at Washington University. The National Chimpanzee Brain Resource is supported by NIH grant NS092988.

## MATERIALS AND METHODS

### Participants

#### Humans

30 human participants (19 female; 11 male; ages between 22 and 36) randomly selected from the database provided by the Human Connectome Project (HCP): https://www.humanconnectome.org/study/hcp-young-adult. This sample has been used previously in studies of LPFC sulcal morphology (Miller et al., 2021a,b). HCP consortium data were previously acquired using protocols approved by the Washington University Institutional Review Board. As our previous morphological analyses of LPFC sulci did not show any sex differences across a range of participant ages (from 6–36; Miller et al., 2021a, Voorhies et al., 2021), we did not specifically balance sex when selecting participants. Additionally, the chimpanzee sample also contains a similar ratio of female to male participants.

#### Chimpanzees

Anatomical T1 scans were previously acquired using MRI in 60 chimpanzees (38 female; 22 male; ages between 9 and 54), and no new data were collected for the present study. 30 chimpanzees were used to create a species-specific average template and were not included in any other analyses. Of the remaining chimpanzees, 29 are included in the manual labeling and morphological analyses. 1 chimpanzee was excluded for substantial errors in the cortical surface reconstruction. These participants have also been used in a previous study of sulcal morphology in ventral temporal cortex (Miller et al., 2020). The chimpanzees were all members of the colony housed at the Yerkes National Primate Research Center (YNPRC) of Emory University. All methods were carried out in accordance with YNPRC and Emory University’s Institutional Animal Care and Use Committee (IACUC) guidelines. Institutional approval was obtained prior to the onset of data collection. Chimpanzee MRIs were obtained from a data archive of scans collected prior to the 2015 implementation of U.S. Fish and Wildlife Service and National Institutes of Health regulations governing research with chimpanzees. These scans were made available through the National Chimpanzee Brain Resource (https://www.chimpanzeebrain.org; supported by NIH grant NS092988).

### Data Aquisition

#### Humans

Anatomical T1-weighted MRI scans (0.8 mm voxel resolution) were obtained in native space from the HCP database, along with outputs from the HCP modified FreeSurfer pipeline.

#### Chimpanzees

Detailed descriptions of the scanning parameters have been described in Keller et al. 2009, but we also describe the methods briefly here. Specifically, T1-weighted magnetization-prepared rapid-acquisition gradient echo (MPRAGE) MR images were obtained using a Siemens 3 T Trio MR system (TR = 2300 ms, TE = 4.4 ms, TI = 1100 ms, flip angle = 8, FOV = 200 mm) at YNPRC in Atlanta, Georgia. Before reconstructing the cortical surface, the T1 of each chimpanzee was scaled to the size of the human brain. As described in Hopkins et al. 2017, within FSL, the BET function was used to automatically strip away the skull, (2) the FAST function was used to correct for intensity variations due to magnetic susceptibility artifacts and radio frequency field inhomogeneities (i.e., bias field correction), and (3) the FLIRT function was used to normalize the isolated brain to the MNI152 template brain using a 7 degree of freedom transformation (i.e., three translations, three rotations, and one uniform scaling), which preserved the shape of individual brains. Next, each T1 was segmented using FreeSurfer. The fact that the brains are already isolated, both bias-field correction and size-normalization, greatly assisted in segmenting the chimpanzee brain in FreeSurfer. Furthermore, the initial use of FSL also has the specific benefit, as mentioned above, of enabling the individual brains to be spatially normalized with preserved brain shape, and the values of this transformation matrix and the scaling factor were saved for later use.

### Manual sulcal labeling

#### Humans

We first manually defined the LPFC sulci within each individual hemisphere in tksurfer (Miller et al., 2021a). Manual lines were drawn on the inflated cortical surface to define sulci based on the most recent schematics of sulcal patterning in LPFC by Petrides (2019), as well as by the pial and smoothwm surfaces of each individual (Miller et al., 2021a, 2021b). In some cases, the precise start or end point of a sulcus can be difficult to determine on a surface (Borne et al., 2020). Thus, using the inflated, pial, and smoothwm surfaces of each individual to inform our labeling allowed us to form a consensus across surfaces and clearly determine each sulcal boundary. For each hemisphere, the location of LPFC sulci was confirmed by trained independent raters (C.B.H., W.I.V., J.A.M.) and finalized by a neuroanatomist (K.S.W.).

#### Chimpanzees

Probability maps generated from the human sulcal labels (Miller et al., 2021a) were used to guide sulcal labeling in chimpanzees (Figure 1a). Binarized maps for posterior middle frontal tertiary sulci (pmfs-p, pmfs-i, pmfs-a) were projected into individual chimpanzee hemispheres as a combined label with freesurfer’s *mris_label2label function*. Additionally, three dorsal and ventral bounding sulci were included in the analysis: The posterior and anterior components of the superior frontal sulcus (sfs_p, sfs_a) and the inferior frontal sulcus (ifs). These human predictions were used to inform the manual identification of these sulci in chimpanzees (Figure 1b). For each hemisphere, we identified between one and three pmfs components. These sulci were labeled posterior, intermediate, or anterior based on their position relative to the bounding sulci as in our previous work (Miller et al., 2021a; Petrides 2019). Tertiary sulci that fell outside of the human pmfs prediction and anterior to the bounding sulci were defined as components of the paraintermediate frontal sulcus (pimfs; Voorhies et al., 2021; Petrides 2019). The pimfs was only present in one chimpanzee (c19). As with humans, for each hemisphere, the location of LPFC sulci was confirmed by trained independent raters (C.B.H., W.I.V., N.S., J.K.Y., C.M.) and finalized by a neuroanatomist (K.S.W.).

### Characterization of sulcal patterning

We characterized the frequency of occurrence of each sulcus separately for left and right hemispheres. We compared the frequency of occurrence of the tertiary sulcal components between hemispheres and species with chi-square tests. For each sulcus, we also characterized sulcal patterns, or types, based on intersections with surrounding sulci. For each sulcal pair, we report the number of intersections relative to the total frequency of occurrence of that sulcus in the hemisphere (Figure 2c). We report Pearson correlation coefficients between left and right hemispheres in each sample, as well as the correlation between species.

### Sulcal morphology

#### Depth

Depth of each sulcus was calculated in millimeters from each native cortical surface reconstruction. Raw values for sulcal depth were calculated from the sulcal fundus to the smoothed outer pial surface using a modified version of a recent algorithm for robust morphological statistics which builds on the Freesurfer pipeline. The original algorithm samples the 100 deepest vertices to determine the fundal depth. To address differences in sulcal size across species and sulci, particularly in the small tertiary sulci, we modified the algorithm to sample the deepest 10% of vertices in a given sulcus. Results are consistent when titrating the percentage of the deepest vertices included in the analyses (Supplementary Figure 2). As the chimpanzee surfaces were scaled prior to reconstruction, we also report relative depth values for the sulci of interest. For these metrics, within each species, depth was calculated relative to the deepest point in the inferior frontal sulcus.

#### Surface area

Surface area (in square millimeters) was generated for each sulcus from the *mris_anatomical_stats* function in FreeSurfer (Dale et al., 1999; Fischl et al., 1999a). Again, to address scaling concerns between species, we also report surface area relative to the total surface area of the frontal lobe.

### Morphological comparisons

All comparisons were conducted using mixed effects linear models implemented in the nlme R package. For both depth and surface area analyses, model predictors included sulcus, hemisphere, and species, as well as their interaction terms. Species, hemisphere, and sulcus were considered fixed effects. Sulcus was nested within hemisphere which was nested within subjects. Post-hoc analyses were computed with the *emmeans* function.

### Asymmetry analyses

For each label, hemispheric asymmetry was computed with the following calculation:

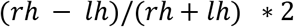

Asymmetry values were computed for each species separately. A linear mixed effects model was used to assess the sulcus by species interaction.

### Probability maps

Sulcal probability maps were calculated to summarize those vertices that had the highest and lowest correspondence across individual chimpanzees, respectively. To generate these maps, each sulcal label was transformed from the individual to a chimpanzee template surface from a held-out population of 30 chimpanzee brains that was made with the FreeSurfer *make_average_subject* function (Miller et al., 2020). Once transformed to this common template space, for each vertex, we calculated the proportion of chimpanzees for whom the vertex is labeled as the given sulcus. In the case of multiple labels, we employed a greedy, “winner-take-all” approach such that the sulcus with the highest overlap across participants was assigned to a given vertex. Consistent with previous studies (Miller et al., 2020; Voorhies et al., 2021) in addition to providing unthresholded maps, we also constrain these maps to maximum probability maps (MPMs) with 20% participant overlap. MPMs help to avoid overlapping sulci and increase interpretability (Figure 5a).

### Data availability

Data and analysis pipelines used for this project will be made freely available on GitHub upon publication. Requests for further information should be directed to the Corresponding Author, Kevin Weiner.

