## Supplementary Figures for "Defining tertiary sulci in lateral prefrontal cortex in chimpanzees using human predictions"

**Supplementary Materials**

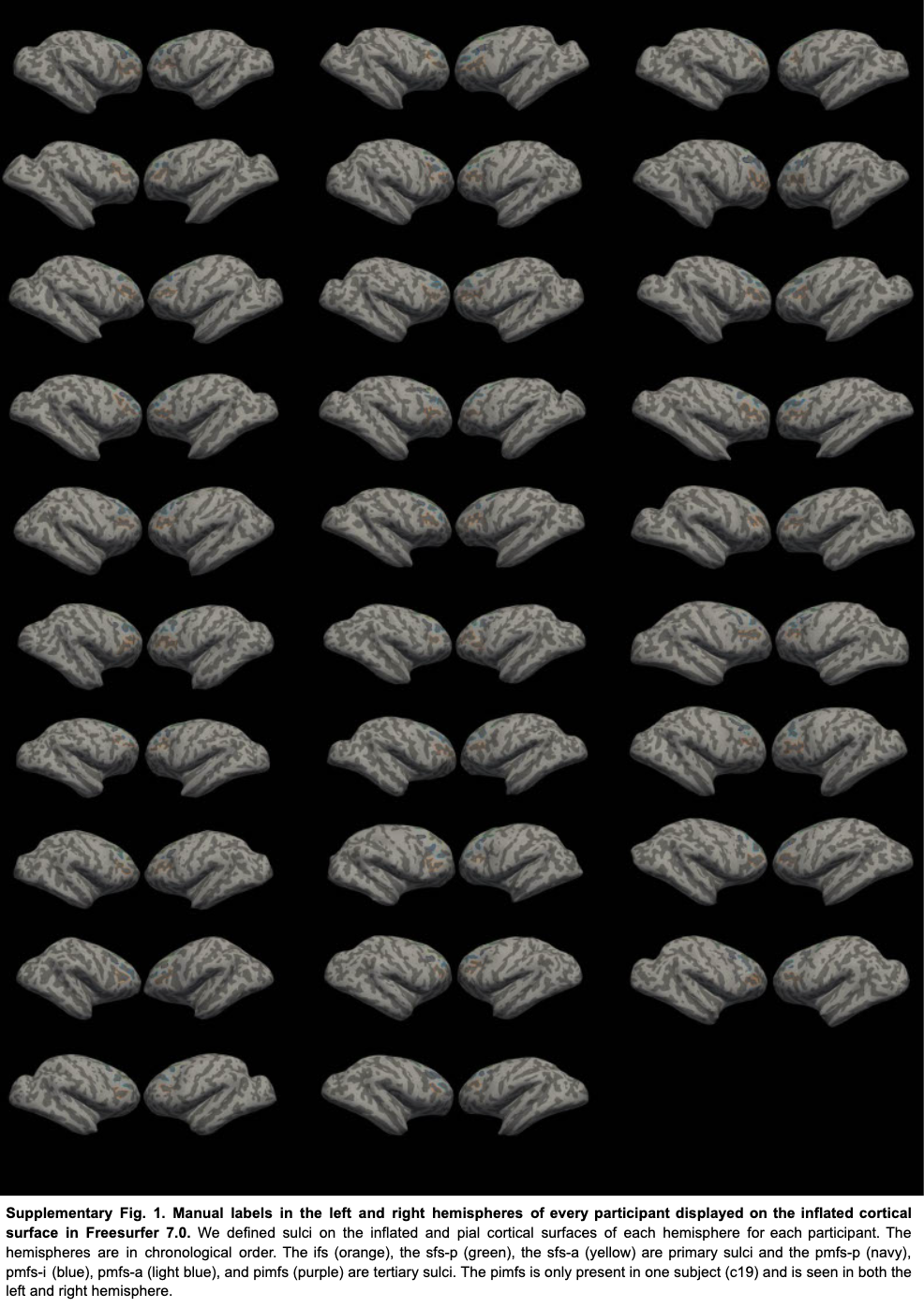

***Supplementary Figure 1. Manual labeling protocol of LPFC tertiary in chimpanzees guided by human predictions.*** We defined sulci on the inflated and pial cortical surfaces of each hemisphere for each chimpanzee. The ifs (orange), sfs-p (dark green), sfs-a (light green), pmfs-p (dark blue), pmfs-i (blue), and pmfs-a (light blue) are consistently identifiable across hemispheres. The pimfs (purple) is only present for one chimpanzee (c19) in the left and right hemisphere.

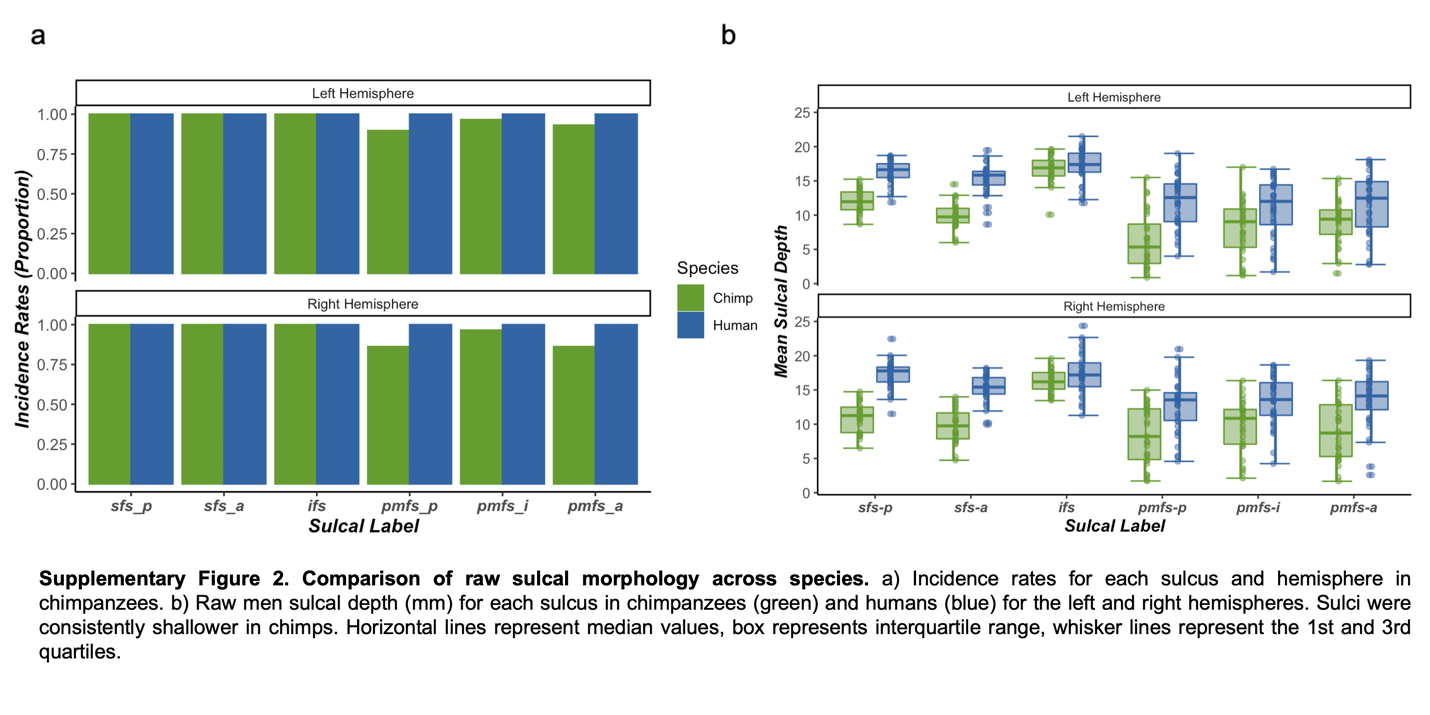

**Supplementary Figure 2. Comparison of raw sulcal morphology across species.** a) Comparison of sulcal incidence rates for chimpanzees and humans. b) Raw mean sulcal depth (mm) for each sulcus in chimpanzees (green) and humans (blue) for the left and right hemispheres. Sulci were consistently shallower in chimpanzees. Horizontal lines represent median values, boxes represent interquartile range, and whisker lines represent the 1st and 3rd quartiles.
